# Nematic structures contribute to robust zygotic polarization in *C. elegans*

**DOI:** 10.64898/2026.03.30.715403

**Authors:** Michiel Vanslambrouck, Jef Vangheel, Ella Linxia Müller, Bart Smeets, Pierre Gönczy, Rob Jelier

## Abstract

The *C. elegans* zygote is a powerful model for asymmetric cell division. Its strikingly patterned cortex features thick F-actin bundles and myosin foci, contractile nematic structures that drive characteristic surface ruffles. Early in the cell cycle, symmetry breaks near the sperm-contributed centrosomes, typically at the presumptive posterior pole, and is marked by local downregulation of contractility. This initiates a cortical flow that polarizes the cell and enables PAR proteins to establish anterior and posterior domains. While biochemical mechanisms maintaining these domains are well understood, the mechanical role of cortical architecture in polarization remains unclear.

We developed a three-dimensional (3D) mechanical model of *C. elegans* zygote polarization that represents the actin bundles and myosin foci of the cortex as a network of stiff contractile filaments. We measured cortical flow with high spatiotemporal resolution by tracking myosin foci, and used these data alongside mechanical properties from the literature to parametrize the model. The model simulates the complete polarization process in 3D, from symmetry breaking through domain stabilization, and reproduces key cortical dynamics including flow profiles, surface ruffles, and tension anisotropy. Domain arrest near the embryo midpoint emerges from density-dependent contractility regulation, in which cortical material redistribution during flow creates a mechanical negative feedback that balances anterior and posterior tension.We find that compressive flow aligns actin bundles in the anterior domain and generates anisotropic tension perpendicular to the flow direction. Although this alignment is not essential for polarization when symmetry breaking occurs at the pole, it contributes to this process when symmetry breaking occurs laterally. In such cases, anisotropic tension from aligned bundles drives axis convergence by rotating the posterior domain towards the nearest pole. Nematic cortical structures therefore ensure robust alignment of the polarization axis.

**Availability:** All data and code required to reproduce the results are freely available at https://doi.org/10.5281/zenodo.18135771. The latest version of the software is maintained at https://bitbucket.org/pgmsembryogenesis/polarization.

## Introduction

Asymmetric divisions are a hallmark of embryogenesis and play a central role in cell differentiation. The zygote of the nematode *Caenorhabditis elegans* is an established model for asymmetric division. Prior to division, the actomyosin cortex exhibits a characteristic polarizing flow that establishes the anterior-posterior (AP) axis. The actomyosin cortex is a dynamic contractile network located immediately beneath the plasma membrane, composed primarily of actin filaments and non-muscle myosin motor proteins (NMY-2, hereafter myosin) and has a major role in determining cell shape (Lecuit et al., 2011).

During the transition from the oocyte to the zygote, cortical tension increases as myosin is activated by the small GTPase RhoA (RHO-1 in *C. elegans*) (Yan et al., 2022; Reymann et al., 2016; Cowan and Hyman, 2007). The resulting highly contractile cortex forms a pattern with transient myosin clusters (which we will refer to as foci for simplicity) connected by thick bundles of actin filaments (Nishikawa et al., 2017) (fig. 1A). These bundles are nematic structures, whereby actin filaments show long-range orientational alignment without strict positional order (Balasubramaniam et al., 2022). Foci are evenly distributed across the cortex initially and the connecting bundles undergo periodic contractions, forming a tensioned network that generates local invaginations or ruffles on the cell surface (Mayer et al., 2010; Munro et al., 2004; Gan and Motegi, 2021) (fig. 1A). Polarization of the zygote begins with a symmetry-breaking cue: the sperm-contributed centrosomes interact with the cortex, which correlates with a local drop in myosin contractility, usually at the presumptive posterior pole of the embryo (fig. 1B) (Albertson, 1984; Goldstein and Hird, 1996; Sadler and Shakes, 2000), as reviewed in (Cowan and Hyman, 2007; Rose and Gönczy, 2014). The region with reduced contractility then expands to form a less dense cortical domain, with dense cortical material flowing towards the opposite, presumptive anterior side, until it covers half of the embryo. During this polarization process, the anterior domain has more myosin than the posterior one, and cortical ruffling is present only anteriorly (Hird and White, 1993; Munro and Bowerman, 2009). Simultaneously, a cytoplasmic counter-flow moves material from the future anterior towards the future posterior (Munro et al., 2004).

**Fig 1.**
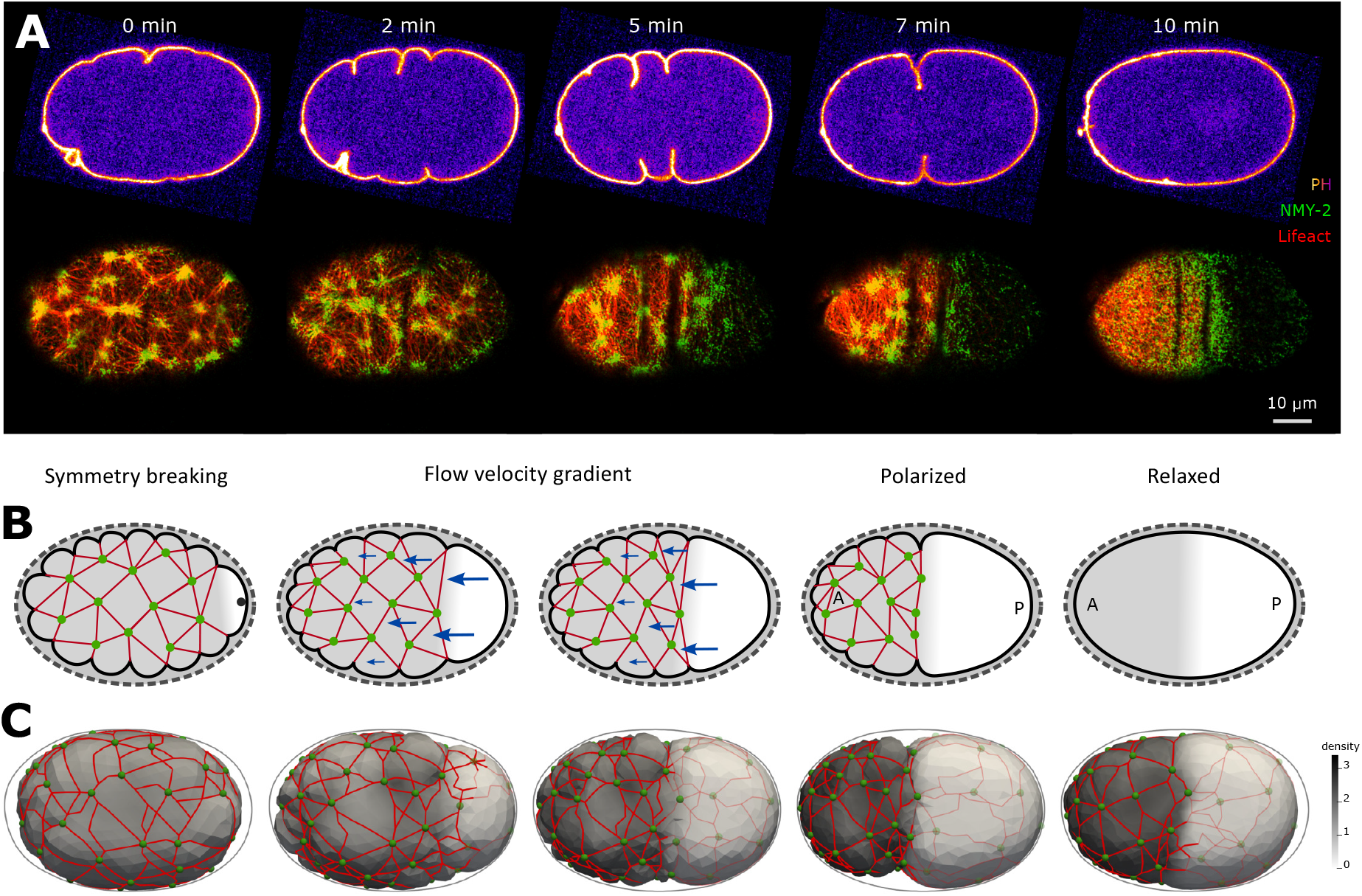
Stages of zygote polarization. A: Fluorescence microscopy of the polarizing zygote. Top panels show plasma membrane (GFP::PH) with clearly visible surface ruffles. Bottom panels show cortical myosin (NMY-2::GFP) and F-actin (Lifeact:mKate2). Nematic structures are formed by bundles of actin filaments, with myosin foci at their junctions. The embryo polarizes, after which large nematic structures disappear and the cortex relaxes. Full time-lapses are provided as Supplementary Videos 1–2. Note that the zygote maintains stable domains beyond the last frame shown (10’), until division, which occurs 15–18’ after symmetry breaking. B: Schematic representation of the different stages of polarization. Initially, the zygote has a uniform actomyosin-rich cortex with a tensile network of actin filament bundles and myosin foci that generate membrane ruffles. Sperm-contributed centrosomes break symmetry at the presumptive posterior pole, which is accompanied by local downregulation of myosin and decreased cortical tension. Thereafter, the actomyosin-rich cortex starts to flow away from this point towards the anterior, and an expanding region with a less dense cortex forms in its wake. Flow continues until the actomyosin-rich cortex covers approximately half of the embryo. At this point, about 7 minutes after symmetry breaking, a pronounced ruffle termed the pseudocleavage furrow forms at the domain boundary. Shortly afterwards, cortical tension decreases, and nematic structures disappear, including ruffles, the pseudocleavage furrow, and myosin foci. C: Simulated cortex model with actin bundles (red) and myosin foci (green) replicating the stages of polarization observed in vivo.

Several mechanical features of the polarizing flow have been uncovered. One early hypothesis stated that cortical flow begins with acute rupture of a prestressed cortex (Munro and Bowerman, 2009). However, the flow does not resemble an elastic relaxation. Instead, cortical flow relies on continuous actomyosin activity: the flow persists for several minutes, far longer than cortical turnover times, demonstrating that this is an actively driven viscous process rather than the release of stored elastic stress (Munro et al., 2004; Munro and Bowerman, 2009; Mayer et al., 2010). Cortical laser ablation experiments during polarization indicated that the cortex has a large hydrodynamic length, the characteristic length of stress propagation (14 µm) (Mayer et al., 2010; Saha et al., 2016). Moreover, ablations revealed tension anisotropy in the cortex, with anterior tension perpendicular to the AP axis twice as high as in the posterior, and tension parallel to the AP axis approximately uniform throughout the zygote (fig. 3A) (Mayer et al., 2010). Mayer et al. (2010) used a continuum viscous fluid model to interpret these observations and argued that cortical viscosity is considerably larger than the friction acting on a moving cortex. In this view, the relatively high viscosity enables the cortex to sustain long-distance flow, with transport of cortical material over tens of micrometers during polarization (Mayer et al., 2010). They also interpreted the tension anisotropy in the anterior cortex to be a consequence of the flows’ compressive effect on the cortex and resulting viscous stresses. However, whether nematic organization of actin bundles contributes to force transmission and mechanical anisotropy of the cortex remains unclear.

The polarizing flow establishes cortical asymmetry, which is then stabilized by biochemical feedback from the PAR proteins (Munro, 2006). Anterior PAR proteins (PAR-3, PAR-6, PKC-3) are localized to the dense anterior cortex, while posterior PAR proteins (PAR-1, PAR-2, LGL-1) are recruited to the nascent, expanding posterior cortex (Cowan and Hyman, 2007; Hoege et al., 2010; Beatty et al., 2010). Domain establishment takes 5–8 minutes following symmetry breaking, after which polarity is maintained for approximately 10 minutes until the first mitotic division (Blanchoud et al., 2015; Oswald, 2010; Munro and Bowerman, 2009; Cowan and Hyman, 2007; Klinkert et al., 2019). During the maintenance phase, a deep surface invagination termed the pseudocleavage furrow forms at the center of the embryo, before receding rapidly (Strome, 1986; Munro et al., 2004). Cortical flow and ruffles cease at the same time as the pseudocleavage disappears (Hird and White, 1993).

In rare cases, the sperm-derived centrosomes are positioned laterally rather than at the pole at the time of symmetry breaking. In such cases, cortical flow is initiated laterally, after which the centrosomes and the smooth cortical domain move to the nearest pole, which then adopts posterior features (Goldstein and Hird, 1996; Reymann et al., 2016; Bhatnagar et al., 2023). This process, termed *axis convergence* (Bhatnagar et al., 2023), reorients the AP axis to align with the egg’s long axis, after which development proceeds normally.

Several models of polarization have been developed using reaction-advection-diffusion equations to describe PAR protein dynamics and domain interactions (Goehring et al., 2011; Blanchoud et al., 2015; Gross et al., 2018). Different studies focused on active gel theory to model cortical mechanics (Reymann et al., 2016; Naganathan et al., 2018; Salbreux et al., 2022; Bhatnagar et al., 2023). In Bhatnagar et al. (2023), the cortex is represented as a compressible thin film with spatially varying myosin concentration, and with actin filaments exhibiting nematic alignment. The distribution of myosin generates tension gradients, which produce flow that aligns filaments into the pseudocleavage furrow, thereby creating anisotropic tension in that location. The authors predict that the pseudocleavage furrow contributes to axis convergence by minimizing its circumference through reorientation perpendicular to the egg’s long axis, which is consistent with several experimental observations (Bhatnagar et al., 2023).

While some studies, such as Bhatnagar et al. (2023), have incorporated nematic structure, they rely on continuum frameworks that spatially average the mechanical properties of the cortex. Consequently, these models do not account for discrete actin bundles, and their influence on flow dynamics and polarization remains unknown. Here we develop a 3D biomechanical cell simulation that explicitly represents actin bundles as tension-generating filaments, to investigate how nematic structures affect cortical flow and polarization. Specifically, we measure cortical flow velocities over time and use them alongside mechanical properties from the literature to parametrize the model and reproduce wild-type polarization dynamics. We then evaluate the role of actin bundles in generating cortical tension and in flow dynamics during axis convergence following lateral symmetry breaking.

## Results

### Model of the cortex

To investigate the mechanical role of nematic structures in zygotic polarization, we extended the Deformable Cell Model (DCM) (Odenthal et al., 2013; Smeets et al., 2019; Cuvelier et al., 2023; Vanslambrouck et al., 2024) to explicitly represent cortical actin bundles and myosin foci. Simulations start with a uniform distribution of cortical material. Symmetry is broken at a defined location by a local drop in tension, after which cortical flow arises spontaneously with a velocity profile comparable to what is observed in vivo. The flow is tuned to arrest when the domain boundary reaches the middle of the embryo, and a stable configuration of anterior and posterior domains is achieved with a size ratio of approximately 1 : 1.

In the DCM, a cell is represented as a triangular mesh with surface tension (*γ*) and viscosity (*η*). Here, we added nematic structures as a network of actin bundles, with myosin foci at their intersections. Foci are initialized randomly on the zygote surface, and neighboring foci are linked by actin bundles. Each bundle forms a chain of consecutive mesh edges connecting two foci, with a local, directed tension *γ*_*L*_ along the chain (fig. 1C). The contractile bundles resist bending, compression, and extension, and thus act as structural components that contribute to the long-range force transmission observed in vivo (Saha et al., 2016).

Using a simple model with a uniform surface tension and a stable mesh of contractile actin bundles, we could observe cortical flows and ruffles. However, the unpolarized zygotic state was unstable: a transient uneven distribution of bundles would locally reduce tension and trigger spontaneous symmetry breaking and ultimately collapse of the contractile network (Supplementary Video 4), reminiscent of spontaneous symmetry breaking observed in vivo in the absence of sperm-derived centrosomal cues (Klinkert et al., 2019; Zhao et al., 2019; Reich et al., 2019; Kapoor and Kotak, 2019). However, the model did not reflect the generally stable pre-polarization zygotic cortex observed in healthy embryos. Since the zygotic cortex is constantly remodeled, with structural turnover of both myosin foci and actin bundles (Munro et al., 2004; Mayer et al., 2010; Saha et al., 2016), we reasoned that introducing such turnover in the model could stabilize the bundle network, and implemented that foci approaching each other merge, while new foci are generated when they move apart (see Materials and Methods). For simplicity, we do not model the variable tension from focal contractions that are observed approximately every minute during polarization (Munro and Bowerman, 2009; Nishikawa et al., 2017), nor do we model chiral cortical flows (Naganathan et al., 2018).

We next partly parametrize the system using data from various publications (full list in Supplementary Table S1). Following Mayer et al. (2010), we assume high cortical viscosity relative to cytoplasmic friction and ignore elastic effects. Atomic force microscopy measurements (Yamamoto et al., 2023) determined surface tension values up to 0.5 nN/µm, which we set as the maximum *γ* in our model. Laser ablation experiments in the anterior cortex showed an initial recoil velocity of approximately 10 µm/min (Mayer et al., 2010; Vanslambrouck et al., 2024). This velocity reflects the ratio of tension to damping coefficient, assuming the cortex behaves as a Kelvin-Voigt material (Mayer et al., 2010). We therefore set bundle line tension and damping coefficient to match this ratio: *γ*_*L*_*/c*_*N*_ = 10 µm/min. Mayer et al. (2010) estimated an inter-foci distance of around 5 µm; we use comparable values with *d*_min_ = 4.5 µm and *d*_max_ = 9 µm.

### Domain stabilization

When the flow comes to a halt with the domain boundary in the center of the zygote in vivo, ruffling is still present in the anterior domain (Hird and White, 1993), and hence the actin bundles still produce significant local contractility. Nonetheless, if flows have ceased at that time, we can infer that both anterior and posterior domains generate similar tension in the AP direction. This happens despite asymmetric distribution of myosin (Supplementary Figure S2), which drives the flows initially, yet remains present as the flows stop, indicating regulation of some kind.

To halt cortical flow in simulations, we introduce density-dependent tension regulation. We track a cortical material density factor *ρ*, initially uniform, that is advected by the flow and increases in the compressing anterior while decreasing in the expanding posterior. This density serves as a proxy for the overall cortical state, including the accumulation of regulatory proteins such as PAR proteins (Gross et al., 2018). We define three phenomenological tension rules, each motivated by experimental observations, described below.

First, we make the bundle line tension *γ*_*L*_ dependent on *ρ*, so that *γ*_*L*_ decreases in regions with low cortex density. Actin bundles require sufficient cortical material to form, and are nearly absent in the posterior region in vivo (Munro et al., 2004). We assume that bundle function is optimal around *ρ* = 1 and decreases linearly to zero with *ρ* (Materials and Methods, fig. 5).

Second, we lower bundle tension in high density domains. Mayer et al. (2010) suggest a saturating relationship between myosin accumulation and contractility, as the densest myosin foci could stall parts of the network upon maximal contraction. Additionally, the E-cadherin HMR-1 was found to accumulate in the anterior cortex together with actin, where it downregulates RHO-1, inhibiting myosin activity (Padmanabhan et al., 2017). Gross et al. (2018) also found that the anterior PAR protein PAR-6 decreases the dissociation rate of myosin, which could affect contractility in the anterior, and shows that PAR proteins are regulating cortical properties. These observations lead us to decrease actin bundle line tension *γ*_*L*_ as the cortex compresses (Materials and Methods, fig. 5).

However, even if actin bundles in the anterior produce less contractile force as polarization nears completion, the posterior cortex has few bundles and foci to counteract forces from the anterior domain. The ablation experiments from Mayer et al. (2010) confirm that the posterior domain, with sparse bundles and foci, matches the anterior’s tension along the AP axis, with similar recoil velocities (fig. 3A). Similar effects have been observed after the division of the EMS cell, where the sparse cortex of E produces more tension than the dense cortex of MS (Caroti et al., 2021; Vanslambrouck et al., 2024). For these reasons, we add a third form of tension regulation in which surface tension *γ* increases as *ρ* decreases over the density range of our simulations (Materials and Methods, fig. 5). This negative feedback counters expansion and compression and stabilizes the system.

### Parameter tuning using flow measurements

We now evaluate the polarization model against experimental observations. We measured cortical flow velocities in control zygotes using fluorescently labeled myosin and utrophin (NMY-2::RFP; GFP::utrophin) Utrophin was chosen as a marker for F-actin filaments instead of LifeAct, as the latter may misrepresent actin bundles by stabilizing filaments (Hirani et al., 2019). Myosin foci were manually tracked over time to obtain high-resolution flow profiles (see Materials and Methods). A smooth spatiotemporal flow profile fit to data from all embryos analyzed is shown in fig. 2A (see Materials and Methods). Shortly after symmetry breaking, the posterior-to-anterior flow arises with a steep speed gradient, reaching average flow rates of 4.3 µm/min from 150 s to 200 s (across the AP axis) while peaking at 9.6 µm/min near the posterior pole before declining. These values are in line with those of the literature, where typical reported speeds are 5 µm/min (Hird and White, 1993; Gross et al., 2018; Blanchoud et al., 2015; Cheeks et al., 2004), up to 7.7 µm/min (Munro et al., 2004). Flow remains slower in the anterior throughout polarization. Flow largely ceases by 400 s when polarization completes, though local flow of around 3 µm/min persists just posterior of the embryo midpoint, suggesting a modest tension gradient across the domain boundary.

**Fig 2.**
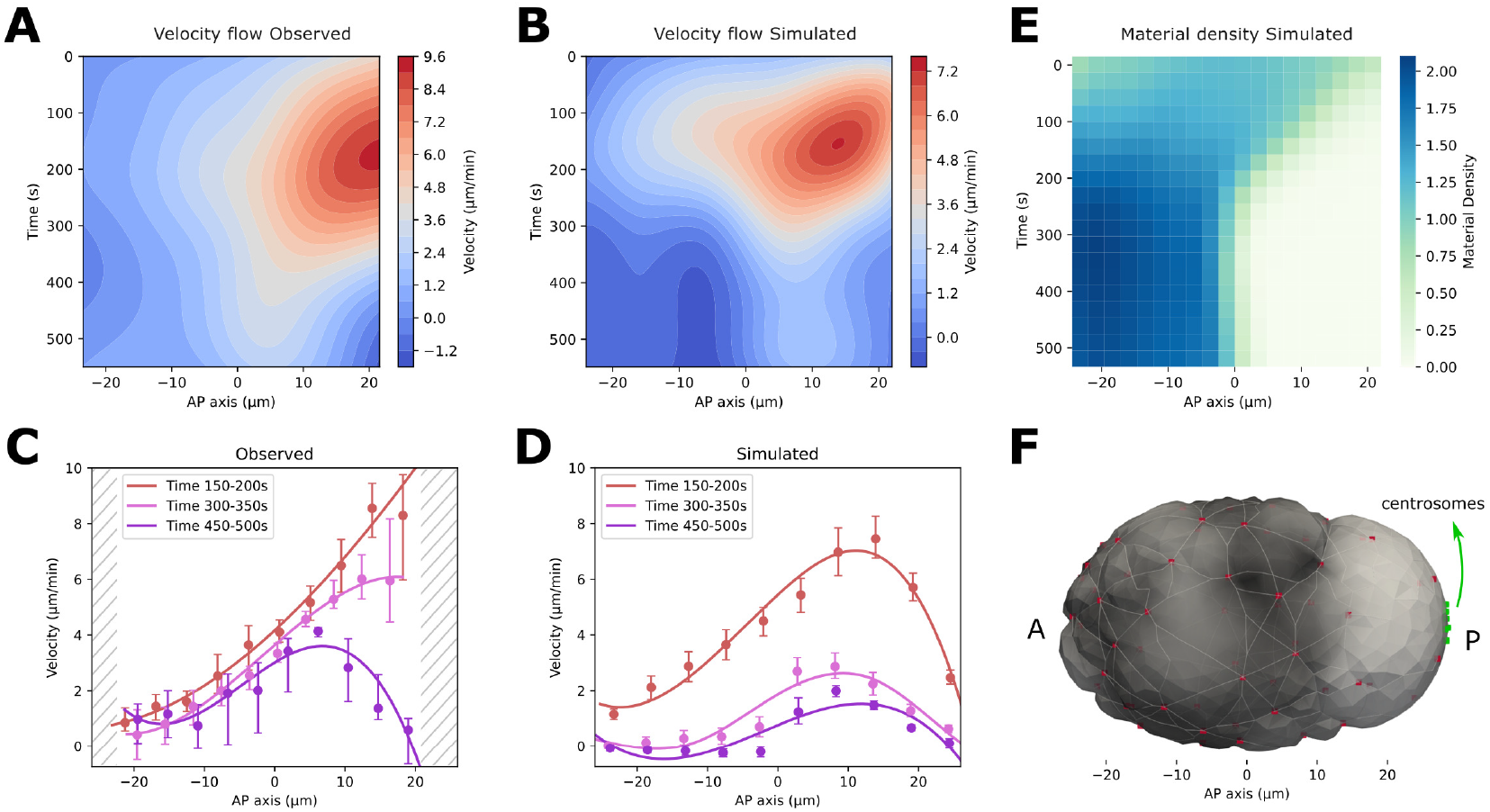
Polarization flow velocity comparison. A, B: Kymographs of flow speed projected on the AP axis, starting at symmetry breaking. Cortical flow velocity measured in vivo (left) from 6 embryos and through simulation (right) with 20 repeats. Note that the color scales differ to highlight similarity in flow patterns; quantitative comparison is found in panels C and D. C, D: Flow velocity graphs with 95% CI (bootstrap, n=1000) markers for several time windows. The cortical imaging setup does not capture the curved, polar-most regions of the embryo (striped regions indicate missing data). E: Kymograph of material density during simulated flow. F: Rendering of simulated embryo for spatial reference. Centrosomes (in green) show where the flow originates and the cortex becomes smooth. Ruffles persist only in the contractile anterior region.

**Fig 3.**
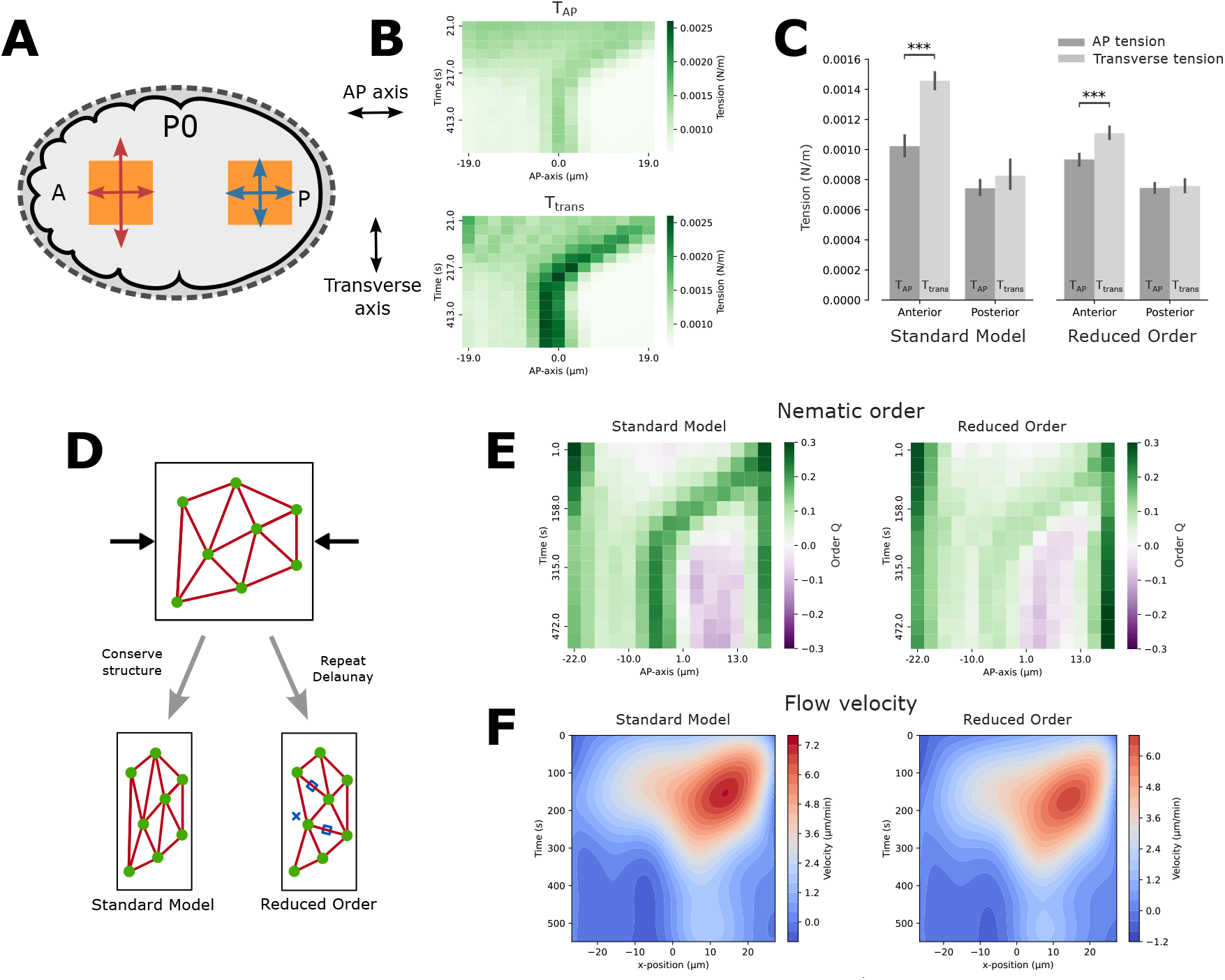
Anisotropic tension and nematic order in simulation. A: Schematic of anisotropic tension in the polarizing zygote derived from laser ablation experiments Mayer et al. (2010). B: Kymographs of directional tension, parallel to AP and transverse axis respectively. C: Anisotropic tension measured in anterior and posterior domain for two cases. Both windows span 180–300 s and cover the transverse axis 12 µm wide. On the AP axis, windows cover regions −13 µm to −1 µm, and 5 µm to 17 µm for anterior and posterior respectively. D: Schematic of process to eliminate nematic order in the bundle network, yielding the ‘Reduced Order’ derived model. E: Kymographs of nematic order *Q* for standard model and Reduced Order case. The central dark green band represents the area that is being compressed most, which shifts from the posterior (right) side to the center of the zygote as the polarization process goes on. The dark bands on the sides show high nematic order because the surface at the poles curves until it is perpendicular to the AP axis. F: Cortical flow velocity for standard model and Reduced Order case.

We fine-tuned model parameters by fitting simulated flow to the experimental flow profiles. Bundle line tension *γ*_*L*_ = 4.5 nN was chosen to halt polarization near the embryo midpoint. Significantly lower values result in incomplete polarization, while higher values drive the domain boundary too far anteriorly (Supplementary Table S2). Polarization speed is primarily controlled by cortical viscosity, which we chose at *η* = 90 kPas to match observed flow speeds and duration (fig. 2B).

Symmetry breaking parameters were also tuned to match experimental observations. Local tension downregulation in the model must exceed a threshold to reliably trigger symmetry breaking (Supplementary Table S2), consistent with experimental findings (Munro and Bowerman, 2009). By default, we initialize the centrosome at the posterior pole, pinned on the cortex, where it can move along with the surface. From the centrosome contact site (fig. 2F), actin bundles continuously break within 4 µm, and experience reduced tension in the surrounding area. This setting reliably triggers symmetry breaking, whereas reducing the break radius delays symmetry breaking.

With this parametrization, we reach the polarization behavior shown in fig. 1C and animated in Supplementary Video 5. To quantify the agreement between simulations and experiments, we compared key metrics of cortical flow dynamics. The simulations reproduced the rapid onset of cortical flow and the AP speed gradient observed experimentally, with peak velocities of 7.5 µm/min occurring 160 s after flow initiation near the posterior pole. The material density (*ρ*) evolution can be followed in fig. 2E, describing how cortical material moves and accumulates in the anterior. The AP domain boundary reaches the midpoint at 280 s in the simulation as in the actual embryo. Moreover, the subsequent decay in flow speed followed a similar time course in both simulations and experiments (fig. 2A, B), with maintenance flows stabilizing around 2 µm/min. Furthermore, the domain boundary remains stable at the midpoint once established (fig. 2E).

Some differences remain between simulated and experimental flow profiles, however (Supplementary Figure S1). The simulated speed gradient is less steep than in the actual embryo during peak flow, and does not reach the highest velocities observed in the posterior domain in vivo (fig. 2C-D). Additionally, the simulated flow decays more rapidly near the posterior pole than in vivo (fig. 2A-B). In the actual embryo, cortical material appears to be recruited or reassembled more efficiently to sustain flow, resulting in a more gradual decline in flow speed compared to simulations (see Discussion).

To assess the robustness of our model, we performed a sensitivity analysis to determine which parameters are critical for maintaining the observed velocity profile and polarized final state. Our simulations produced stable polarization across a wide range of parameter values (Supplementary Table S2), which shows that the model does not depend on a specific parameter set and is consistent with the well-established robustness of *C. elegans* zygote polarization in vivo. Certain parameters, particularly those governing actin bundle density and bundle line tension, affected the rate of polarization (Supplementary Table S2). However, the final ratio between anterior and posterior domains remained remarkably consistent (0.45–0.54 across all parameter sets). This robustness emerges naturally from the coupling between contractility and cortical flows in our model (contractility causes flow, which changes *ρ*, which regulates contractility), suggesting that such coupling is sufficient to explain the system’s inherent resilience to perturbations.

### Anisotropic tension and bundle alignment

We next examine whether the model reproduces the anisotropic cortical tension observed experimentally, and what structural mechanism underlies it. Cortical laser ablation experiments by Mayer et al. (2010) showed that during polarization, when the anterior domain still covers 70% of the embryo, anterior cortical tension in the AP direction is approximately half of that in the transverse direction (fig. 3A). In the posterior, by contrast, AP tension in both directions is similar, so that no anisotropy is present. Figure 3B shows directional tension (*T* in our model) over time along the AP and transverse directions. We find that the simulated tension profile is consistent with values inferred from the laser ablation experiments. Transverse tension *T*_*trans*_ changes most over time and space, being elevated in the anterior, and peaking at the domain boundary upon polarization completion. AP tension shows a similar spatiotemporal pattern but remains weaker in the anterior and changes less in magnitude. Restricting the tension values to the spatiotemporal window of zygote ablations (fig. 3C) confirms that *T*_*trans*_ exceeds *T*_*AP*_ (*p <* 10^−7^, standard model). Since the model only includes uniform surface tension and contractile actin bundles, bundle alignment is the cause of this tension increase.

Can actin bundles actually align? Although the turnover of F-actin and myosin molecules at this stage of development is rapid compared to the flow duration (10–30 s) (Mayer et al., 2010; Saha et al., 2016; Robin et al., 2014), the larger bundles and resulting surface ruffles can last for several minutes. Munro et al. (2004) measured myosin focus lifetimes around 120 s, so foci can persist through a large part of polarization. Our cortex model incorporates limited structural turnover, with filament bundles persisting on a timescale of minutes. As the cortex flows to the anterior, it is continuously compressed, which allows actin bundles to align. This parallel alignment can be described as nematic order.

To computationally explore the contribution of bundle alignment to the polarization process, we derived a version of the model in which any alignment that arises is immediately destroyed, followed by rebuilding of the bundle network (Reduced Order, fig. 3D). To quantify bundle alignment, we use a statistic for nematic order *Q* similar to Reymann et al. (2016), where every bundle’s angle with the AP axis is used to measure directionality bias in the network (Materials and Methods, eq. 1). When *Q* is negative, bundles align along the AP axis, and when it is positive, bundles align in the transverse direction. In fig. 3E, a band of high nematic order (dark green) forms near the posterior pole at polarization onset and shifts toward the center as polarization progresses. The anterior region maintains slightly positive nematic order due to compression, whereas the posterior region shows slightly negative order due to expansion. The Reduced Order model (fig. 3E, right) confirms that most compression-induced nematic order is eliminated by rebuilding of the network, though some remains. Further, in fig. 3C, we see that limiting nematic order decreases anterior tension in the transverse direction, though the anisotropy between *T*_*AP*_ and *T*_*trans*_ remains significant (*p <* 10^−5^).

To assess the effect on polarization, we compare velocity kymographs for both models in fig. 3F and observe that polarization flow is largely unaffected by the reduction of nematic order. These results suggest that the strong anisotropy in the anterior domain arises from flow-induced compression and bundle alignment, but that this anisotropy is not necessary for cortical flow or domain formation.

### Nematic order and axis convergence

Since anisotropic tension from actin bundle alignment is not required for the polarizing flow when sperm-contributed centrosomes are located at the presumptive posterior pole, we investigated whether it contributes to robust polarization under other conditions. Therefore, we tested whether nematic order from thick actin bundles facilitates alignment of AP polarization with the egg’s long axis when symmetry breaking occurs laterally, away from a pole (Goldstein and Hird, 1996; Reymann et al., 2016; Bhatnagar et al., 2023). Figure 4A shows such an event, which occurs occasionally in control embryos. In such a case, the embryo polarizes and rotates to realign the AP axis, with NMY-2 (cyan) concentrating in the anterior and PAR-2 (magenta) in the posterior. We replicate this event in simulations by positioning the symmetry-breaking site laterally (fig. 4B).

**Fig 4.**
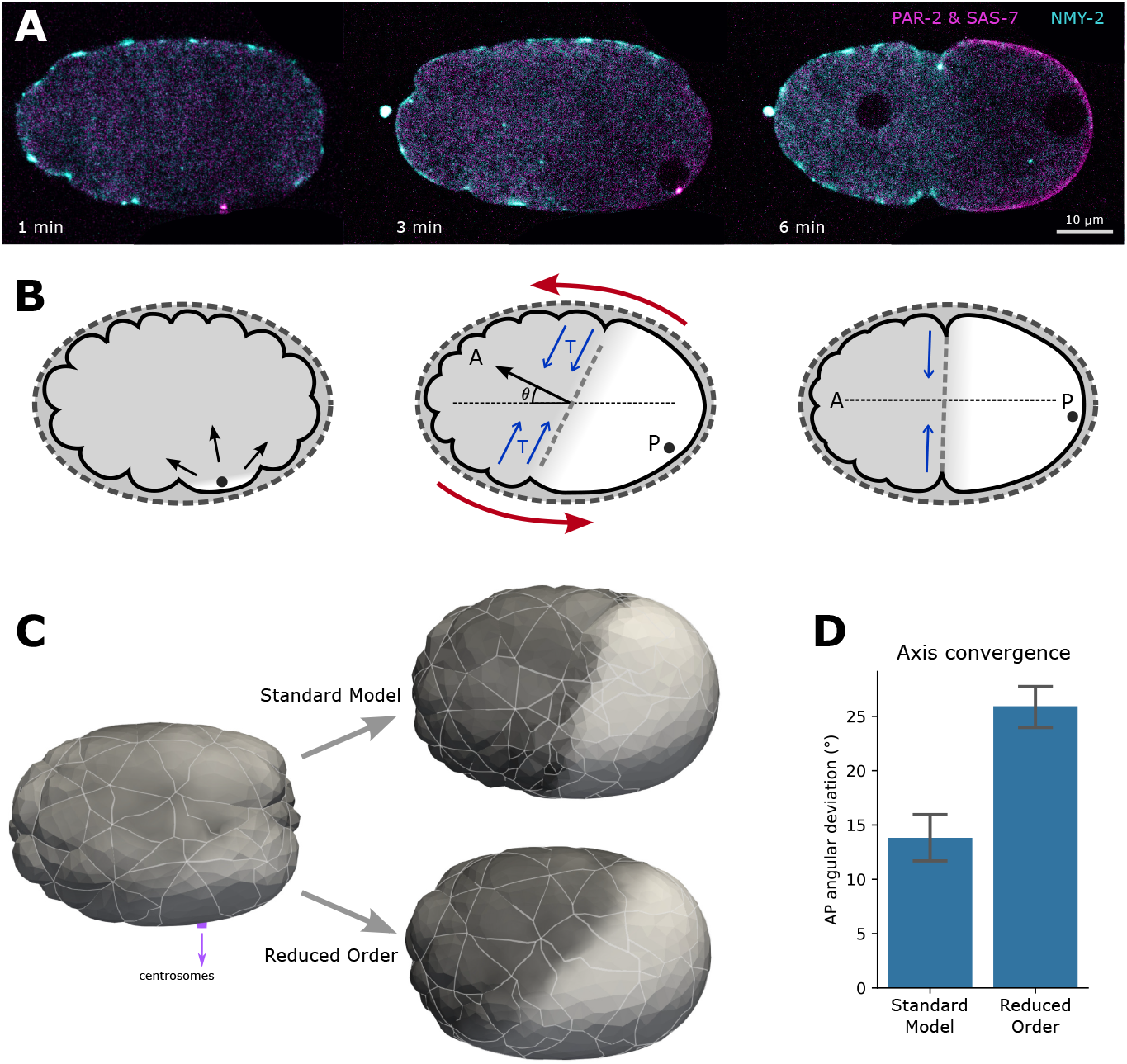
Axis convergence and the effects of nematic order. A: Axis convergence experiment with lateral induction and alignment of the AP axis with the long axis of the egg. PAR-2 and SAS-7 in magenta, NMY-2 in cyan. Full time-lapse is provided as Supplementary Video 3. B: Schematic of lateral symmetry breaking, tension buildup perpendicular to the flow, and axis convergence. C: Axis convergence error *θ* with and without nematic order in the bundle network. Bootstrap (n=1000) was used to calculate 95% confidence intervals.

**Fig 5.**
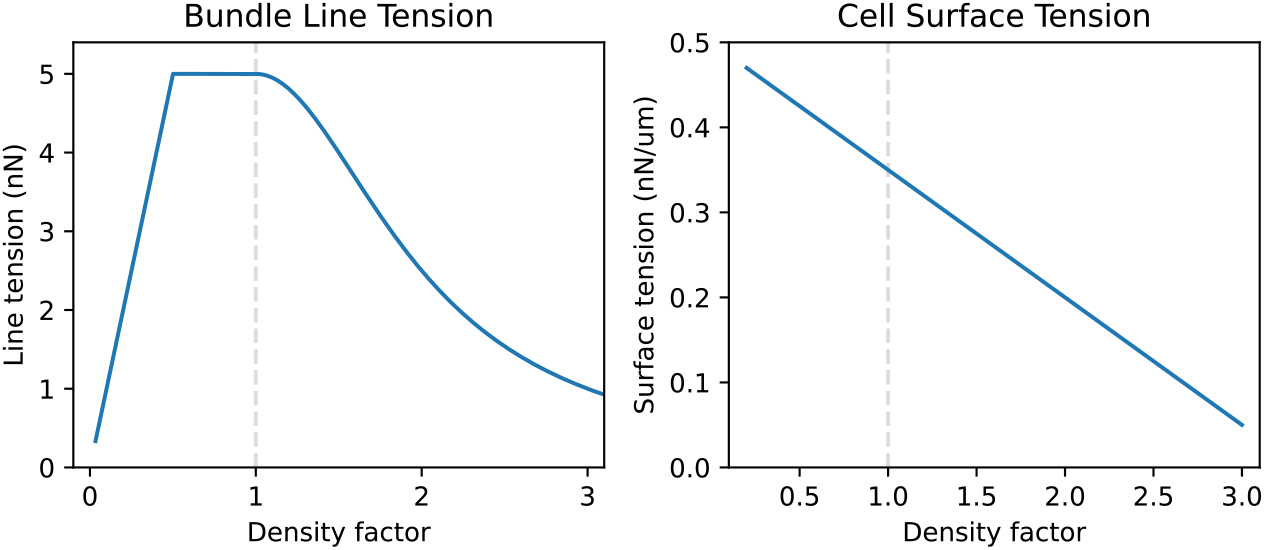
Tension forces in function of cortical material density. A: Line tension for bundles builds up linearly, is maximal around density factor 1, and then decreases quadratically. B: Cortex surface tension decreases linearly in function of density.

We find that such simulated embryos exhibit spontaneous migration of the centrosomes and posterior domain toward the nearest pole (Supplementary Video 6), matching experimental observations. To quantify the process, we use the angle between the axis of polarization (drawn through the center of the domains) 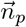 and the egg’s long axis as the angular axis error. We find that axis convergence is slower in simulations than in vivo. Since this dynamic process depends on multiple parameters beyond those constrained by flow dynamics (see section *Parameter tuning using flow measurements*), we extend simulations to 1100 s after polarization induction to allow convergence to complete. With this adjustment we find an axis error of 14^◦^ in the standard model (fig. 4C-D). To assess the effect of nematic order, we remove bundle alignment as in the previous section. This adjustment increases the axis error to 26^◦^, significantly higher than in the standard model (*p* = 3 *×* 10^−10^). We conclude that nematic order and the resulting tension anisotropy are necessary for axis convergence upon lateral symmetry breaking, and therefore contribute to robust polarization of the zygote.

## Discussion

We developed a 3D mechanical model of *C. elegans* zygote polarization that represents cortical nematic structures (actin bundles and myosin foci) as a network of stiff contractile filaments. The model reproduces the stable prepolarization stage with a remodeling network that generates regular cortical ruffles. Symmetry breaking initiates polarizing cortical flow through a local reduction in cortical tension at the centrosome contact site. We quantified cortical flow by tracking myosin foci in control zygotes to create high-resolution spatiotemporal profiles. These flow data, combined with mechanical properties from the literature, allowed us to parametrize the model and reproduce wild-type polarization dynamics. Under the compressive flow in the anterior domain, actin bundles align and generate nematic order. This alignment produces the observed tension anisotropy but is not necessary for flow or domain formation. However, nematic order plays a role in axis convergence: when symmetry breaking occurs laterally, anisotropic tension from aligned bundles drives posterior domain rotation toward the nearest pole. Nematic cortical structures therefore contribute to robust polarization in C. elegans.

Under compressive flows, the actin bundle network can build up nematic order, yet this order does not influence the dynamics of the polarizing flow itself. In simulation, we observe strong bundle alignment in the central region of the embryo, at the domain boundary, where the pseudocleavage also forms later in vivo, consistent with previous observations (Reymann et al., 2016; Bhatnagar et al., 2023). Our simulations also show a gradual buildup of alignment across the entire anterior side during compression, which generates anisotropic tension. This result matches experimental observations of anisotropic tension in the anterior domain (Mayer et al., 2010), but suggests a different mechanism. Mayer et al. (2010) attributed tension reduction along the compression axis to internal viscosity resisting compressive flow, assuming uniform contractility and without actin bundles being explicitly represented. However, posterior ablations show isotropic tension (Mayer et al., 2010) despite velocity gradients being present (fig. 2A and C, 150–200 s), and ablation velocity decays along the AP axis are similar across domains. Our model demonstrates that bundle alignment can generate anterior anisotropy without requiring spatially varying material properties, providing a structural explanation consistent with the observed cortical network organization.

Our results show that actin bundle alignment is sufficient to correct laterally induced polarization. This correction is driven by anisotropic tension that orients the contractile domain boundary to minimize its circumference, positioning it perpendicular to the embryo’s long axis. Bhatnagar et al. (2023) proposed that axis convergence results from flow-driven actin alignment interacting with embryo geometry. In their model, the cortex is treated as an active nematic fluid layer in which a myosin-dependent contractility gradient, set by posterior centrioles, drives cortical and cytoplasmic flows that reposition the posterior pronucleus. Nematic effects arise from a homogeneous population of actin filaments that align under flow, without explicit cortical transport of myosin or other cortical proteins. Our interpretation is distinct in two ways. First, individual actin filaments turn over rapidly (10–30 s) (Mayer et al., 2010; Saha et al., 2016; Robin et al., 2014), which limits persistent alignment by flow. Instead, the cortex contains pre-existing nematic structures in the form of actin bundles with associated myosin foci that persist for minutes (Munro et al., 2004) and are present before polarization. We suggest that compressive flow aligns these stable bundles, and that their alignment generates the anisotropic tension that drives axis convergence. Second, Bhatnagar et al. (2023) attributed axis convergence primarily to the pseudocleavage furrow. However, axis convergence occurs during active polarization flow, whereas the pseudocleavage furrow appears near the end of polarization (400 s after symmetry breaking). Our simulations suggest that anisotropic tension from progressive bundle alignment throughout polarization drives convergence, with the pseudocleavage furrow marking the peak of this alignment at the domain boundary rather than initiating it. Consistent with this interpretation, we frequently observed spontaneous formation of a deep furrow at the center of the zygote near the end of polarization in our simulations (fig. 1C), matching the observed timing and position of the pseudocleavage. This supports the hypothesis proposed by Reymann et al. (2016) that pseudocleavage arises as a consequence of strong nematic order generated by compressive flows.

Cortical flow nearly stops around 400 s after symmetry breaking. Although anterior ruffling persists, which shows that actin bundles still contract, this flow cessation implies that tension generated by anterior and posterior domains is balanced at this stage. In our model, this balance arises through a negative feedback loop: flow redistributes cortical material, which locally changes how tension is generated. Specifically, the increase in cortical density in the compressing anterior reduces actin bundle line tension. This is consistent with anterior accumulation of cortex-associated proteins that downregulate myosin activity, such as E-cadherin. Conversely, the posterior expands, is highly dynamic and has lower actomyosin density, yet generates relatively higher surface tension in our model. Such an apparently counterintuitive relationship between lower protein levels and higher tension was previously observed in the E cell of the *C. elegans* embryo (Caroti et al., 2021; Vanslambrouck et al., 2024). One potential explanation is that the posterior domain has a more optimal level of connectivity at intermediate density, as was shown in mitotic HeLa cells where peak tension occurs at intermediate cortical thickness (Chugh et al., 2017). Through this opposing response, tension differences diminish as polarization proceeds. The sensitivity analysis (Supplementary Table S2) shows that although kinetic parameters alter polarization rate, the domain sizes remain roughly equal. This stability follows from the density-dependent contractility regulation. We therefore propose that cortical material redistribution during flow provides a mechanical negative feedback that balances anterior and posterior tensions prior to biochemical stabilization by PAR proteins. Here, cortical density represents overall cortical state changes rather than literal material amounts, a hypothesis that requires experimental validation.

Our simulations reproduce key features of cortical flow but show quantitative discrepancies with experimental velocity profiles, particularly in peak flow speeds and flow persistence. These differences likely reflect biophysical properties absent from our minimal model, including spatially varying cortical viscosity, heterogeneous turnover rates, or nonlinear feedback between flow and contractility. We deliberately kept model complexity to the minimum sufficient to reproduce zygotic cortex behavior, adding only the actin bundle network and density-dependent contractility regulation to the DCM. Spatial variations in cortical viscosity, contractility, or friction could affect the exact flow dynamics, but these factors are largely unknown, as measurements are lacking and difficult to obtain. We therefore chose not to introduce additional free parameters to describe such phenomena. The model is grounded in experimental measurements from the literature, and we characterized cortical flow at high spatiotemporal resolution to identify the key features the simulations must reproduce.

Several extensions could improve the model’s realism. Axis convergence simulations reveal incomplete capture of viscous forces, which suggests the DCM model may need to be refined. Retrograde cytoplasmic flow and focal contractions producing periodic contractility bursts (Munro and Bowerman, 2009) are also not yet included. Our mechanical framework deliberately omitted biochemical dynamics to maintain tractability, but future work could integrate PAR protein association and dissociation kinetics with cortical material density. Such coupling would test whether mechanical domain establishment precedes and influences biochemical polarization, or whether both processes are intrinsically linked. The model also provides a basis to study how nematic structures interact with biochemical polarity cues in other developmental contexts where cortical architecture and biochemical signaling coordinate.

This work provides an explicit three-dimensional modeling of the zygotic cortex, carefully parametrized against experimental observations, and offers insight into how multiple observed phenomena (flow-induced bundle alignment, anisotropic tension, domain stabilization, and axis convergence) combine to complement the primary polarization mechanism. Our mechanical framework provides a foundation for future studies connecting cortical architecture to biochemical domain stabilization and to symmetry-breaking events in other developmental contexts.

## Materials and Methods

### *C. elegans* Strains and maintenance

*C. elegans* strains were grown on NGM plates at 20 ^◦^C as described by Brenner (1974).

For fig. 1A, we used imaging from two strains for membrane and actomyosin respectively:

- GZ1559 (nmy-2(ges6(nmy-2::RFP-T+unc-119(+)))I; ltIs38[pAA1; pie-1::GFP::PH(PLC1delta1) + unc-119(+)]).
- RJ012, a cross of LP162 (nmy-2(cp13(nmy-2::GFP + LoxP)) I.) (Dickinson et al., 2015) and SWG001 (gesls001 Pmex-5::Lifeact:mKate2::nmy-2UTR, unc-119+1) (Reymann et al., 2016).

For focus tracking we used GZ2131 (nmy-2(ges6(nmy-2::RFP-T+unc-119(+)))I; xsSi3 [GFP::utrophin + Cbr-unc-119(+)] II), a cross between SWG008 and MG589.

For lateral symmetry breaking in fig. 4A, we used GZ1476 (nmy-2(ges6(nmy-2::RFP-T+unc-119(+)))I; par-2(it328[gfp::par-2]) sas-7(or1940[gfp::sas-7])III), previously described by Klinkert et al. (2019).

### Imaging

Embryos were dissected from gravid hermaphrodites in Shelton’s growth medium (Shelton and Bowerman, 1996) and mounted with 20 µm polystyrene beads. Dual-color fluorescence time-lapse imaging was conducted on a Visitron CSU W1 spinning disk confocal microscope with a 60× U Plan S-Apo oil objective (NA 1.42) connected to an Orca Flash 4.0 sCMOS camera, using a 488 nm and 561 nm laser. Full z-stacks were required (covering 30 µm in steps of 1 µm) for GZ1559. For lateral symmetry breaking, GZ1476 embryos were imaged every 10 s in the plane where pronuclei and eggshell are in focus. For cortical imaging, GZ2131 embryos were imaged at the cortex every 5 s using a z-stack covering 2 µm around the cortex in steps of 0.5 µm.

RJ012 embryos were mounted on slides with M9 buffer and compressed to 20 µm with Polybead Microspheres (Polysciences). Imaging was conducted using a Zeiss LSM 880 confocal microscope with a Plan-Apochromat 63×/1.4 DIC M27 oil immersion objective.

### Observed flow measurements

Myosin foci of polarizing GZ2131 embryos were manually tracked using in-house tracking software. The series are synchronized with symmetry breaking at time 0. We only use the velocity component parallel to the AP axis. To reduce erratic changes in velocity, tracks were smoothed using a moving average that includes both the previous and the next frame. fig. 2A was made in Python by fitting a 2D tensor spline (package *pygam*, function *LinearGAM*) on all the speed data over time and the AP axis. The fitted spline was plotted using *contourf* from the package *matplotlib*.

### Cell mechanical model

Zygote simulations are performed using Mpacts, a particle-based simulation framework to compute the motion of interacting particles based on a force balance (Odenthal et al., 2013; Smeets et al., 2019). The core model setup is based on previous cell simulation publications (Cuvelier et al., 2023; Ongenae et al., 2024; Vanslambrouck et al., 2024; Vangheel et al., 2025). The cell is modeled as a foam-like material using the DCM, which considers the cell cortex and membrane as a viscous shell under tension. Forces like tension and contact pressure act on the nodes of a triangular mesh, causing realistic deformation of the cell’s shape. A cell is parametrized by cortical thickness *t*_*c*_, surface tension *γ*, and viscosity *η*. It has a reference volume 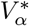, and experiences resistance to volume change controlled by a bulk modulus *K*_*V*_, which exerts additional pressure *P* on the cell membrane. Interactions with the eggshell are managed through repulsion. Validations of the DCM regarding geometry and contact angles are provided in Cuvelier et al. (2023). At the cell-eggshell interface there is also wet friction with friction coefficient *ξ*, while medium viscosity acts on all nodes. We also keep track of a cortical material density factor *ρ*, describing the expansion and compression of triangle area. This density is slowly equalized to neighboring triangles with a diffusive flux 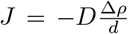, where *D* is the diffusivity and *d* the distance between two triangles. The material transfer between triangles is then dependent on the length of the shared edge *L* and the time step Δ*t*: *T* = *J · L*Δ*t*. A more in-depth description with a force balance per node is included in SI.

### Nematic structure

During polarization, the zygotic cortex forms a network of thick actin bundles and contractile myosin foci. We model this as a separate layer on top of the mesh representing the cortex, connected via the nodes. To initialize this network, we evenly sample the surface to assign cell nodes as foci using Trimesh `sample_surface_even` (Dawson-Haggerty et al.). A 3D Delaunay triangulation on the foci positions and internal scaffold points gives us a way to efficiently connect foci on the surface. On those connections, actin bundles are defined as a surface linker from focus to focus. To integrate them with the rest of the cortex, every bundle follows the shortest path from focus to focus, via the existing triangle edges of the mesh. Tensile forces will then be placed on these edges, forcing the path to become straight.

The network is kept stable by spawning and removing foci based on the distance between them. When two foci get closer than *d*_min_, they get merged and their bundles are combined. They will maintain a bundle to any focus either of them were connected to, but duplicates are removed. When a cell node gets further than *d*_max_ from any focus, it will become a new focus to fill in the gap, and make new bundles to existing foci using the Delaunay triangulation. We also remesh the surface, splitting and combining triangles to maintain a typical triangle size. Both foci and bundles get repaired under remeshing damage to keep them as close to original as possible.

Bundles define a series of connected edges in between two foci, and are modeled as a line tension with a damper. Each edge experiences line tension *γ*_*L*_ to mimic actomyosin contractility, in addition to normal (*c*_*N*_) and tangential (*c*_*T*_) damping to provide a viscous effect. Line tension is considered optimal around *ρ* = 1. In low density areas, line tension gets reduced linearly (fig. 5A) as cortical material gets depleted. Meanwhile, we assume a quadratic downregulation of line tension in high density areas. For *ρ >* 1, line tension is multiplied with 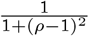 (see Supplementary Information).

Surface tension is also made dependent on the density factor. We assume a negative linear relationship (fig. 5B), where a sparse cortex is highly contractile, while a dense cortex is more stable. In practice, the density in the anterior is close to 2 during simulation. So the dense cortex will still experience a lot of contractile forces from the bundles present, whereas a sparse cortex will not.

Polarization gets induced by the centrosomes, and its location *x*_*induction*_ is pinned on the cell mesh. Under the assumption of cortex disassembly, any bundles within *r*_*induction*_ continuously get removed. Additionally, under the assumption of downregulation, bundles’ tension is reduced to zero at *x*_*induction*_, linearly building up to the normal level at distance 3 *r*_*induction*_.

### Nematic order and anisotropic tension

We measure nematic order *Q* in a manner similar to Reymann et al. (2016), by using every bundle segments length *L* and angle with the AP axis *θ*, to measure a directionality bias of the network (eq. 1).

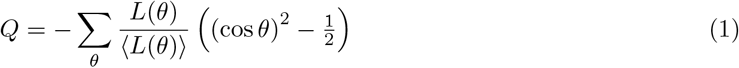

When *Q* is positive, it means there is alignment perpendicular to the AP axis, whereas a negative *Q* means there is parallel alignment.

To measure the directional tension in the simulated cortex, we combine tension information from uniform surface tension and local bundle line tension. In case of the kymographs in fig. 3B, a rectangle patch of 40 µm × 20 µm is evaluated across bins, whereas fig. 3C uses two 12 µm × 12 µm square patches as shown in fig. 3A. Surface tension *T*_*S*_ depends on the local material density and is averaged to find a baseline isotropic tension in nN*/*µm. Anisotropic forces are evaluated in the AP direction (*x*) and the transverse direction (*y*) by using their segment length *L*_*i*_, density dependent line tension *T* (*ρ*_*i*_), and unit orientation 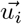

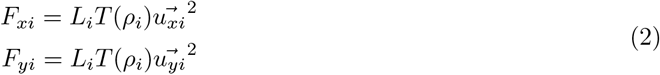

Where we use squared unit vector components so that the total force is correct: 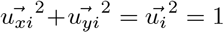. We then add all of the directed bundle forces together and divide by the sampled surface area *S* to get a tension result in nN*/*µm, which can be added to the uniform surface tension to get the total anisotropic tension in direction x and y.

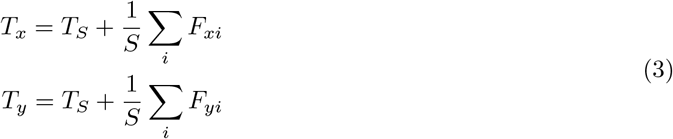

## Acknowledgments

We want to thank Jiř’i Pešek for help with Mpacts and Docker. Additionally, Casper van Bavel, for development of the tracking software and Latex support. Experiment expenses were supported by FWO Research grant number G008423N. M.V. and J.V. were supported by FWO Aspirant grants 1194222N and 11D9923N respectively. E.M. was supported by the Swiss National Science Foundation (grant #310030_197749 to P.G.).

## Notes

https://doi.org/10.5281/zenodo.18135771

### Competing Interest Statement

The authors have declared no competing interest.

