## Supplementary Information for "Nematic structures contribute to robust zygotic polarization in *C. elegans*"

#### 1 Supplementary text

##### 1.1 DCM force balance

Here, we provide the details of the DCM, based on Cuvelier et al. [1]. Cells behave as pressurized liquid "bubbles" with surface tension and internal pressure. The actomyosin cortex of a cell is represented by a viscous shell which is approximated by a triangulated surface mesh with effective thickness $t_c$, and nodal positions $\mathbf{x}_i$. In the following sections we will discuss different force contributions and their implementation.

###### Surface tension

Isotropic tension generated in the cortex results from actomyosin contractility, which is represented as an effective surface tension $\gamma$ in the shell model. Using the derivation of Fedosov et al. [2], the direct force contribution of surface tension $\gamma$ of node $i$ is given by

$$\mathbf{F}_i^{\mathrm{act}} = \frac{\gamma}{2}(\mathbf{x}_k - \mathbf{x}_j) \times \hat{\mathbf{n}}_A, \tag{1}$$

where $i, j, k$ are nodes of a single triangle $A = (ijk)$.

###### Internal pressure

Since the cytoplasm is assumed to be incompressible, active volume control is implemented via a cytoplasmic pressure. We assume the equilibrium volume $V_\alpha^*$ of a cell $\alpha$ is constant and implemented a proportional-integral (PI) pressure controller to enforce this. The cytoplasmic pressure $P_\alpha^{\mathrm{cyt}}(t)$ due to the volume controller is then estimated as

$$P_\alpha^{\mathrm{cyt}}(t) = -K\epsilon_\alpha(t) - \frac{K}{T_I}\int_0^t \mathrm{d}s\, \epsilon_\alpha(s), \tag{2}$$

for a cell with volumetric strain $\epsilon_\alpha(t) = (V_\alpha(t) - V_\alpha^*)/V_\alpha^*$. $T_I$ is typically chosen as one order of magnitude larger than the time-resolution with which the system is updated. The total pressure acting on a node

$$P_i(t) = \frac{2\gamma}{R_\alpha^{\mathrm{cell}}} + P_\alpha^{\mathrm{cyt}}(t),$$

includes an additional offset pressure $2\gamma/R_\alpha^{\mathrm{cell}}$ to balance the actomyosin contractility in the cortex. This ensures cells are in mechanical equilibrium at the start of the simulations. The force acting on node $i$ due to nodal pressure $P_i(t)$, is obtained by integrating over the node associated oriented Voronoi area $S_i$. Given the pressure is constant over the cell surface, the force is simply given by

$$\mathbf{F}_i^{\mathrm{cyt}}(t) = \mathbf{S}_i\, P_i(t). \tag{3}$$

##### Medium damping

A general drag force

$$\mathbf{F}_i^{\text{drag}} = -\Gamma_{ii}^{\text{f}} \cdot \mathbf{v}_i,$$

where

$$\Gamma_{ii}^{\text{f}} = \frac{3\eta^{\text{f}}}{2R^{\text{cell}}} S_i \mathbb{I}, \tag{4}$$

is included to account for the liquid drag between the cells and their medium due to the fluid viscosity $\eta^{\text{f}}$. When dealing with arbitrary shapes this approximation is no longer correct. Even though $F_i^{\text{drag}}$ is small compared to other dissipative forces, it is still used to improve the stability of the numerical integration scheme as it ensures the resistance matrix is positive definite, see further in equation of motion. [3]

##### Cortex viscosity

Actomyosin remodeling is modeled as an effective cortex viscosity $\eta_c$ as per Vangheel et al. [4]. We distinguish between in-plane and out-of-plane bending viscosity. The in-plane viscous damping force between two nodes $i, j$ with velocity $\mathbf{v}$ is given by

$$\mathbf{F}_i^{\eta_c} = \frac{\eta_c t_c}{\sqrt{3}} \Big( (\hat{\mathbf{n}}_{ij} \cdot (\mathbf{v}_j - \mathbf{v}_i))\hat{\mathbf{n}}_{ij} + (\hat{\mathbf{t}}_{ij} \cdot (\mathbf{v}_j - \mathbf{v}_i))\hat{\mathbf{t}}_{ij} \Big),$$

where the direction of viscous forces are given by the unit vectors

$$\hat{\mathbf{n}}_{ij} = \frac{\mathbf{v}_j - \mathbf{x}_i}{\|\mathbf{v}_j - \mathbf{x}_i\|},$$

$$\hat{\mathbf{t}}_{ij} = \frac{\hat{\mathbf{n}}_A + \hat{\mathbf{n}}_B}{\|\hat{\mathbf{n}}_A + \hat{\mathbf{n}}_B\|} \times \hat{\mathbf{n}}_{ij},$$

with $\hat{\mathbf{n}}_A$, and $\hat{\mathbf{n}}_B$, the normals of the triangles making up the segment between nodes $i$ and $j$. Note that we do not distinguish between bulk, extensional ($\eta_e$), and shear viscosity ($\eta_c$), effectively setting the Trouton ratio $\eta_e/\eta_c$ to 1. In contrast, a Newtonian fluid typically has a Trouton ratio of 3.

The viscous bending moment between two adjacent triangles $A = (i, j, k), B = (i, l, j)$ with small thickness $t_c$ arises from the rate of angle deviation $\frac{\text{d}\phi}{\text{d}t}$ as

$$\|\mathbf{M}_{AB}^{\eta_c}\| = \frac{\eta_c t_c^3}{3} \frac{\text{d}\phi}{\text{d}t}.$$

From this bending moment, we derive the lever forces on the opposing nodes,

$$\mathbf{F}_k^{\eta_c} = -\frac{\mathbf{M}_{AB}^{\eta_c} \times \hat{\mathbf{l}}_A}{l_A},$$

$$\mathbf{F}_l^{\eta_c} = +\frac{\mathbf{M}_{AB}^{\eta_c} \times \hat{\mathbf{l}}_B}{l_B},$$

with $l$ and $\hat{\mathbf{l}}$ the length and unit vector from common axis $(i, j)$ to lever point $k$ or $l$.

For lever $A$, this is

$$\mathbf{l}_A = \hat{\mathbf{n}}_A \times (\mathbf{x}_j - \mathbf{x}_i),$$

$$l_A = (\mathbf{x}_k - \mathbf{x}_i) \cdot \mathbf{l}_A.$$

The forces on the common points $(i, j)$ are weighted with the projected distance of the lever on the common axis,

$$\begin{aligned}
y_{i,A} &= (\mathbf{x}_k - \mathbf{x}_i) \cdot (\mathbf{x}_j - \mathbf{x}_i),\\
y_{j,A} &= (\mathbf{x}_j - \mathbf{x}_k) \cdot (\mathbf{x}_j - \mathbf{x}_i),\\
y_{i,B} &= (\mathbf{x}_l - \mathbf{x}_i) \cdot (\mathbf{x}_j - \mathbf{x}_i),\\
y_{j,B} &= (\mathbf{x}_j - \mathbf{x}_l) \cdot (\mathbf{x}_j - \mathbf{x}_i),
\end{aligned}$$

to obtain the forces on the common nodes for lever forces $F_k^{\eta_c}$ and $F_l^{\eta_c}$

$$\mathbf{F}_i^{\eta_c} = -\frac{y_{j,A}}{y_{i,A} + y_{j,A}} \mathbf{F}_k^{\eta_c} - \frac{y_{j,B}}{y_{i,B} + y_{j,B}} \mathbf{F}_l^{\eta_c},$$

$$\mathbf{F}_j^{\eta_c} = -\frac{y_{i,A}}{y_{j,A} + y_{i,A}} \mathbf{F}_k^{\eta_c} - \frac{y_{i,B}}{y_{i,B} + y_{j,B}} \mathbf{F}_l^{\eta_c},$$

that ensure force balance.

##### Contact forces

The contact pressure on contacting triangles $AB$ is given by

$$P_{AB}^d(\mathbf{x}) = -\frac{k_d}{S_{AB}} \delta(\mathbf{x}), \tag{5}$$

with $k_d$ the contact stiffness, $S_{AB}$ the contact area, and $\delta(\mathbf{x})$ the overlapping distance.

The resulting contact forces and moment are obtained by integrating the total contact pressure over the oriented contact area $\mathbf{S}_{AB}$,

$$\mathbf{F}_{AB}^{\text{adh}} = \int \mathrm{d}\mathbf{S}_{AB}(\mathbf{x})\ P_{AB}^{\text{adh}}(\mathbf{x}). \tag{6}$$

To distribute the contact forces acting on the contact planes to the nodal contact forces $\mathbf{F}_{AB,i}^{\text{adh}}$, it is assumed that the nodal forces are collinear with the contact normal $\hat{\mathbf{n}}_{AB}$. This results in a system of linear equations per contact pair $(AB)$

$$\begin{aligned}
\sum_{i \in A} \mathbf{F}_{AB,i}^{\text{adh}} &= - \int\limits_{\mathbf{x} \in A \cap B} \mathrm{d}\mathbf{S}_{AB}(\mathbf{x})\ P_{AB}^{\text{adh}}(\mathbf{x})\\
\sum_{i \in A} \left[ (\mathbb{I} - \hat{\mathbf{n}}_{AB} \otimes \hat{\mathbf{n}}_{AB}) \cdot (\mathbf{x}_i - \mathbf{x}_{AB}) \right] \times \mathbf{F}_{AB,i}^{\text{adh}} &\\
&= \int\limits_{\mathbf{x} \in A \cap B} \mathrm{d}\mathbf{S}_{AB}(\mathbf{x}) \times (\mathbf{x} - \mathbf{x}_{AB})\ P_{AB}^{\text{adh}}(\mathbf{x})
\end{aligned} \tag{7}$$

The eggshell itself is assumed undeformable, meaning we don't integrate the node positions.

##### Wet contact friction

A viscous contact force is included to account for drag between contacting surfaces with friction constant $\xi$ (units of Pa s/m), expressing the scaling between a dissipative traction $\mathbf{T}_c$ and the sliding velocity of two contacting surfaces $A$ and $B$

$$\mathbf{T}_{AB}^{\text{fric}} = -\xi \Delta \mathbf{v}_{AB}. \tag{8}$$

In the particle-based model, the contact drag force acting on node $i$ of triangle $A$ of the contact pair $(AB)$ is computed as

$$\mathbf{F}_{AB,i}^{\text{fric}} = \Gamma_{AB}^{\text{fric}} \cdot \sum_{k \in B} w_{ik}^{AB} \left(\mathbf{v}_k - \mathbf{v}_i\right) \tag{9}$$

determined by a friction tensor $\Gamma_{AB}^{\text{fric}}$ and weights $w_{ik}^{AB}$ per node $k$ of the $B$ triangle. $w_{ik}^{AB}$ are assumed to scale with the relative contribution of the nodal contact forces to the overall contact force thus

$$w_{ik}^{AB} = \frac{\left(\mathbf{F}_{AB,i}^{\text{adh}} + \mathbf{F}_{AB,k}^{\text{adh}}\right) \cdot \hat{\mathbf{n}}_{AB}}{6 \sum\limits_{\forall k \in B} \mathbf{F}_{AB,k}^{\text{adh}} \cdot \hat{\mathbf{n}}_{AB}}$$

is used as an approximation. $\Gamma_{AB}^{\text{fric}}$ for a given contact area $S_{AB}$ is estimated as

$$\Gamma_{AB}^{\text{fric}} = S_{AB} \left[\xi^{\perp} \hat{\mathbf{n}}_{AB} \otimes \hat{\mathbf{n}}_{AB} + \xi^{\parallel} \left(\mathbb{I} - \hat{\mathbf{n}}_{AB} \otimes \hat{\mathbf{n}}_{AB}\right)\right],$$

with normal and tangential friction coefficients $\xi^{\perp}$ and $\xi^{\parallel}$ respectively. Note that (9) can be equivalently formulated as

$$\mathbf{F}_{AB,i}^{\text{fric}} = \sum_{k \in B} \Gamma_{AB,ik}^{\text{fric}} \cdot \left(\mathbf{v}_k - \mathbf{v}_i\right),$$

if we denote

$$\Gamma_{AB,ik}^{\text{fric}} = w_{ik}^{AB} \Gamma_{AB}^{\text{fric}}. \tag{10}$$

##### Actin bundle forces

Actin bundles define a chain of consecutive edges that link two foci on the mesh. Each chain acts as a contractile element with line tension and a damper that resists bending, compression, and extension. Line tension is regulated by density factor $\rho$ as determined by

$$f(\rho) = \begin{cases} 2\,\rho, & \rho < 0.5, \\ 1, & 0.5 \le \rho \le 1, \\ \dfrac{1}{1 + (\rho - 1)^2}, & \rho > 1. \end{cases}$$

The line tension force on node i, from each connected node j, is

$$\mathbf{F}_{ij}^{\mathrm{L}} = f(\rho)\,\gamma_L\,\hat{\mathbf{n}}_{ij}, \tag{11}$$

With normal direction

$$\hat{\mathbf{n}}_{ij} = \frac{\mathbf{x}_j - \mathbf{x}_i}{\|\mathbf{x}_j - \mathbf{x}_i\|}$$

The damper part of the actin bundles is also affected by density $\rho$, but only in the sparse domain:

$$g(\rho) = \begin{cases} 2\,\rho, & \rho < 0.5, \\ 1, & 0.5 \le \rho. \end{cases}$$

The damping force on node $i$ arises from its relative velocity vector $\mathbf{v}_{ij} = \mathbf{v}_j - \mathbf{v}_i$ with respect to each connected node $j$:

$$\begin{aligned} \mathbf{F}_{ij}^{\mathrm{C}} &= g(\rho)\,c_N\big(\hat{\mathbf{n}}_{ij}\cdot\mathbf{v}_{ij}\big)\hat{\mathbf{n}}_{ij} \\ &\quad + g(\rho)\,c_T\big(\hat{\mathbf{t}}_{ij}\cdot\mathbf{v}_{ij}\big)\hat{\mathbf{t}}_{ij} \\ &\quad + g(\rho)\,c_T\big(\hat{\mathbf{b}}_{ij}\cdot\mathbf{v}_{ij}\big)\hat{\mathbf{b}}_{ij} \end{aligned} \tag{12}$$

Where $c_N$ is the normal damping constant, $c_T$ the tangential damping constant, $\hat{\mathbf{n}}_{ij}$ the unit normal vector from j to i, while $\hat{\mathbf{t}}_{ij}$ and $\hat{\mathbf{b}}_{ij}$ are the two directions perpendicular to $\hat{\mathbf{n}}_{ij}$ so that the three from an orthonormal basis.

##### Equation of motion

In the overdamped cellular environment, inertial forces may be neglected [5]. Based on the different contributions described above, the force balance for node $i$ gives

$$\begin{aligned} &\mathbf{F}_i^{\mathrm{act}} + \mathbf{F}_i^{\mathrm{cyt}} + \sum_j \mathbf{F}_{ij}^{\mathrm{L}} + \sum_{(AB):i\in A} \mathbf{F}_{AB,i}^{\mathrm{adh}} \\ &\qquad = \Gamma_{ii}^{\mathrm{f}}\cdot\mathbf{v}_i + \sum_j \Gamma_{ij}^{\mathrm{c}}\cdot(\mathbf{v}_i - \mathbf{v}_j) + \sum_{(AB):i\in A}\sum_{k\in B}\Gamma_{AB,ik}^{\mathrm{fric}}\cdot(\mathbf{v}_i - \mathbf{v}_k) + \sum_j \mathbf{F}_{ij}^{\mathrm{C}} \end{aligned}$$

which can be abbreviated as

$$\mathbf{F}_i = \sum_j \Gamma_{ij}\cdot\mathbf{v}_j \tag{13}$$

for a system consisting of $N$ nodes, where

$$\Gamma_{ij} = \begin{cases} \Gamma_{ii}^{\mathrm{f}} + \sum_{k:k\neq i}\Gamma_{ik}^{\mathrm{c}} + \sum_{(AB):i\in A}\sum_{k\in B}\Gamma_{AB,ik}^{\mathrm{fric}}, & i = j, \\ -\Gamma_{ij}^{\mathrm{c}} - \sum_{(AB):i\in A, j\in B}\Gamma_{AB,ij}^{\mathrm{fric}}, & i \neq j. \end{cases}$$

The Cartesian components of the overall force vectors can be represented as a single $(3N\times 1)$ column matrix, while the friction matrices can be assembled to a single $(3N\times 3N)$ sparse symmetric positive definite friction matrix [6]. The conjugate gradient method

(CGM) is used to efficiently solve the system for the node velocities $\{\mathbf{v}_j\}$ at each iteration. Positions of the nodes are subsequently updated using a semi-implicit Euler integration scheme.

$$\mathbf{x}_i(t+\Delta t) = \mathbf{x}_i(t) + \Delta t\, \mathbf{v}_i(t+\Delta t), \tag{14}$$

where $\mathbf{v}_i(t+\Delta t)$ are the projected new velocities obtained by solving eq. 13 at time $t$ via the CGM.

#### 2 Supplementary figures

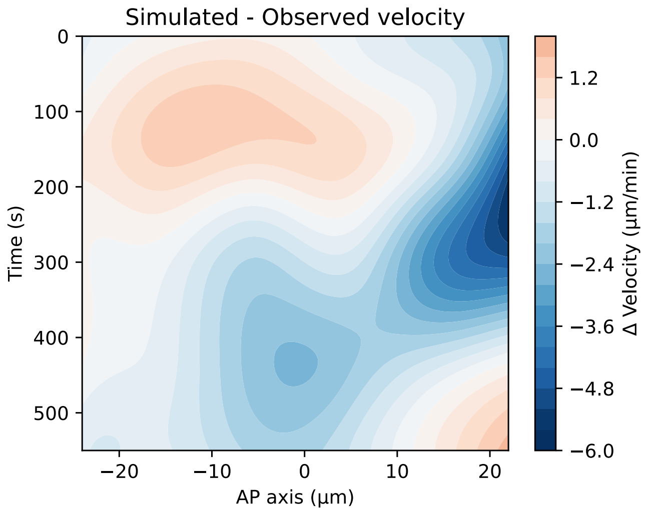


Figure S1: **Kymograph of flow speed difference between simulated (Fig. 2A) and observed (Fig. 2B) polarization.** Biggest error is visible near the posterior side at the end of polarization, as the observed flow perists longer and stronger than simulations could reproduce.

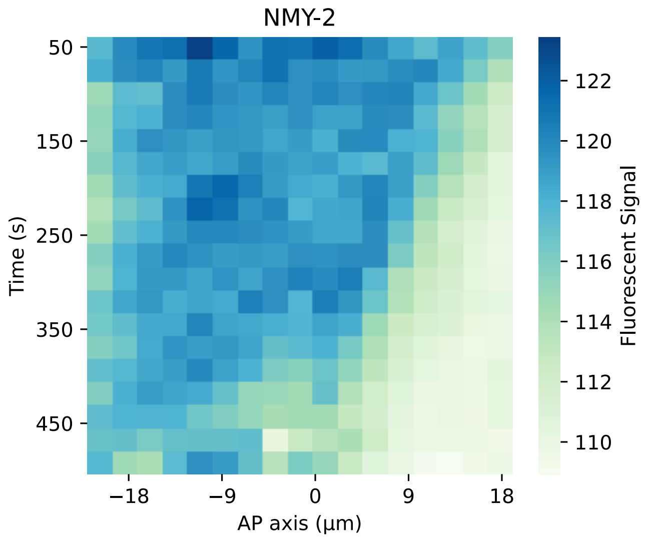


Figure S2: **Kymograph of observed myosin signal during polarization for 6 different embryos.** Myosin moves anteriorly, where the cortical density remains similar to its density before polarization. At the bottom center, a gap is present where the pseudocleavage formed and the fluorescent signal vanished.

#### 3 Supplementary tables

| Parameter | Symbol | Value | Units | Sources |
|---|---|---|---|---|
| Maximum surface tension | $\gamma$ | 0.5 | nN/µm | [7] |
| Local area modulus | $K_A$ | 0.001 | nN/µm | helps with stability |
| Bulk modulus cell | $K_V$ | 1 | kPa | [1] |
| K integral term | $T_I$ | 0.5 | s | |
| Thickness cortex | $t_c$ | 300 | µm | [8, 1] |
| Adhesive range | $h$ | 200 | nm | |
| Normal Friction | $\xi^{\perp}$ | 1 | kPa s/µm | |
| Tangential Friction | $\xi^{\parallel}$ | 0.001 | kPa s/µm | |
| Liquid viscosity | $\eta^{\mathrm{f}}$ | 150 | Pa s | |
| Cortex viscosity | $\eta$ | 90 | kPa s | |
| Diffusivity | $D$ | $2 \times 10^{-14}$ | $\mathrm{m}^2/\mathrm{s}$ | |
| Cell radius | $R_0$ | 19 | µm | [1] |
| Simulation timestep | $\Delta t$ | 0.08 | s | stable |
| Remeshing time | | 1 | s | |
| Bundle line tension | $\gamma_L$ | 4.5 | nN | |
| Normal damping | $c_N$ | 45 | nN s/µm | 10× line tension |
| Tangential damping | $c_T$ | 15 | nN s/µm | |
| Minimum focus distance | $d_{\mathrm{min}}$ | 4.5 | µm | [9] |
| Maximum focus distance | $d_{\mathrm{max}}$ | 9 | µm | |
| Induction range | $r_{induction}$ | 4 | µm | |
| Simulation duration | | 650 | s | |
| Bundles start | | 20 | s | |
| Induction start | | 100 | s | |

Table S1: **Simulation parameters.** Three sections separate continuum properties, particle-based properties, and timings.

| Parameter | Value | $\lambda_p$ | $v_{90}$ | Notes |
|---|---|---|---|---|
| Line tension | 3 | 0.45 | 2.77 | Slow, incomplete polarization, few ruffles |
| $\gamma_L$ (nN) | 4.5 | 0.53 | 4.77 | Default |
| | 6 | 0.54 | 7.62 | Fast polarization, deep ruffles |
| Area conservation | .0005 | 0.54 | 4.82 | – |
| $K_A$ (nN/µm) | .001 | 0.53 | 4.77 | Default |
| | .002 | 0.49 | 4.60 | – |
| Shell friction | .0005 | 0.52 | 4.76 | – |
| $\xi^{\parallel}$ (kPa s/µm) | .001 | 0.53 | 4.77 | Default |
| | .005 | 0.54 | 4.63 | – |
| Cortex viscosity | 80 | 0.54 | 5.01 | Faster polarization |
| $\eta$ (kPa s) | 90 | 0.53 | 4.77 | Default |
| | 100 | 0.53 | 4.49 | Slower polarization |
| Focus distances | 3–7 | 0.58 | 5.92 | Faster due to more bundles |
| $d_{\min}$–$d_{\max}$ (µm) | 4.5–9 | 0.53 | 4.77 | Default |
| | 6–11 | 0.52 | 3.80 | Slower due to less bundles |
| Induction range | 2 | 0.49 | 4.13 | Slower due to weaker induction |
| $r_{induction}$ (µm) | 4 | 0.53 | 4.77 | Default |
| | 6 | 0.54 | 5.09 | Faster due to stronger induction |
| Normal bundle damping | 30 | 0.52 | 4.77 | – |
| $c_N$ (nN s/µm) | 45 | 0.53 | 4.77 | Default |
| | 60 | 0.54 | 4.67 | – |
| Tangential bundle damping | 5 | 0.53 | 4.53 | – |
| $c_T$ (nN s/µm) | 15 | 0.53 | 4.77 | Default |
| | 30 | 0.53 | 4.52 | – |

Table S2: **Sensitivity analysis for physical parameters of polarization simulations.** For each mutation, 10 repeats were simulated. We use two metrics to summarize the polarization flow and resulting domains. To condense the flow speeds into a single number, we use the 90th percentile (0.9 quantile) of the observed focus velocities: $v_{90}$. At the end of the simulation, we use the material density on the cortex mesh to quantify the final domain sizes. First, we find the plane that separates the A and P domains at the 0.1 density quantile. This yields the unit normal vector of this plane $\mathbf{n}_p$ (pointing to A) as the direction of polarization, and its signed distance to the origin $d_p$. We then rescale this to $\lambda_p = (d_p + L/2)/L$ with $L = 52\,\mu\text{m}$ as the typical length of the embryo. This describes a 0–1 range that covers everything from a case without polarization up to a case where the P domain covers the entire zygote. Induction ranges lower than 2 µm experienced occasional delayed or absent symmetry breaking.

### 4 Supplementary videos

**Video 1:** Cortex surface during polarization. Lifeact in red, NMY-2 in green (Frames used in Figure 1).

**Video 2:** Membrane cross-section during polarization. PH marker in LUT (Frames used in Figure 1).

**Video 3:** Polarization with spontaneous axis convergence. PAR-2 and SAS-7 in magenta, nmy-2 in blue (Frames used in Figure 4).

**Video 4:** Incomplete cortex simulation without turnover or tension regulation, leading to network collapse. Foci are rendered in green, actin bundles in red, and density in grayscale.

**Video 5:** Standard polarization simulation with foci in green, actin bundles in red, and density in grayscale.

**Video 6:** Axis convergence simulation with foci in green, actin bundles in red, and density in grayscale. The expanding posterior domain migrates from the lateral induction site to the pole.
